# A comparison of short-read, HiFi long-read, and hybrid strategies for genome-resolved metagenomics

**DOI:** 10.1101/2023.10.04.560907

**Authors:** Raphael Eisenhofer, Joseph Nesme, Luisa Santos-Bay, Adam Koziol, Søren Johannes Sørensen, Antton Alberdi, Ostaizka Aizpurua

## Abstract

Shotgun metagenomics enables the reconstruction of complex microbial communities at a high level of detail. Such an approach can be conducted using both short-read and long-read sequencing data, as well as a combination of both. To assess the pros and cons of these different approaches, we used 22 faecal DNA extracts collected weekly for 11 weeks from two respective lab mice to study seven performance metrics over four combinations of sequencing depth and technology: i) 20 Gbp of Illumina short-read data, ii) 40 Gbp of short-read data, iii) 20 Gbp of PacBio HiFi long-read data, and iv) 40 Gbp of hybrid (20 Gbp of short-read + 20 Gbp of long-read) data. No strategy was best for all metrics, but instead, each one excelled across different metrics. The long-read approach yielded the best assembly statistics, with the highest N50 and lowest number of contigs. The 40 Gbp short-read approach yielded the highest number of refined bins. Finally, the hybrid approach yielded the longest assemblies, and the highest mapping rate to the bacterial genomes. Our results suggest that while long-read sequencing significantly improves the quality of reconstructed bacterial genomes, it is more expensive and requires deeper sequencing than short-read approaches to recover a comparable amount of reconstructed genomes. The most optimal strategy is study-specific, and depends on how researchers assess the tradeoff between the quantity and quality of recovered genomes.

**Importance:** Our understanding of microbial communities is limited by the technologies we employ. Here, we test several different DNA sequencing techniques to better understand the pros and cons of each. Long read DNA sequencing allowed for the reconstruction of higher quality and even complete microbial genomes, however, the cost was greater than commonly used short-read DNA sequencing. We suggest researchers consider the trade-offs between each method and decide based on the goals of their research question/s.

## Introduction

Shotgun metagenomics is increasingly becoming the preferred approach for characterising the genetic composition of complex microbial communities, both in environmental and host-associated samples (1). This is because, unlike targeted amplicon sequencing approaches, shotgun metagenomics enables the reconstruction of microbial genomes through metagenomic assembly and binning, making it a powerful tool (2–4). This process yields metagenome-assembled genomes (MAGs) (5), which can be annotated to provide an overview of the functional capabilities of each genome (6).

DNA sequencing technologies employed for generating shotgun metagenomic data can be grouped in two main categories (7). Short-read sequencing technology, which is the main strategy currently employed, yields large amounts of sequence data at a low cost. However, the short length of generated sequences (usually up to 300 bp) can hinder the assembly of microbial genomes, as the underlying assembly graphs lack the information required to deal with repetitive and duplicated sequences (8). Long-read sequencing technologies can help overcome such limitations. While originally limited by high sequence error rates, novel strategies such as the PacBio HiFi sequencing enable the recovery of sequence qualities comparable to short reads (9). PacBio HiFi has recently been applied to various sample types, including human and sheep faecal samples (10, 11), chicken intestinal samples (12), anaerobic digesters (13), seawater (14), and saline lake sediments (15). However, the costs per data unit for long-read technologies are still several orders of magnitude higher than short-read technologies, which has hampered their widespread adoption. Hybrid assembly, which combines the strengths of both short- and long-read data, has been proposed as a cost-efficient strategy that enables taking advantage of the benefits of both approaches (16–18).

To the best of our knowledge, only one study has compared the quality of MAG assembly from short-read to PacBio HiFi sequencing (15). Tao *et al.* found that HiFi assembly provided the most complete bacterial and viral genomes, and biosynthetic gene cluster, while a hybrid approach yielded a higher quantity and quality of MAGs. In this study, we compare the use of short-read only, long-read only, and hybrid sequencing strategies for recovering MAG catalogues from the faecal microbial communities of laboratory mice. We measured performance by means of multiple metrics targeting quantitative and qualitative traits of the reconstructed MAG catalogues. Based on our results, we discuss the suitability of using different sequencing strategies in light of study design, scientific aims, and available resources.

## Methods

### Animal experiments and sample collection

The experiment was conducted at the ZIBA Experimentation Centre, Zarautz (Basque Country), between September and December 2020, under the animal experimentation licence PRO-AE-SS-154 issued by the Regional Government of Gipuzkoa. The mice sampled for this study were part of a larger project. Diversity Outbred mice of both sexes were acquired from the Jackson Laboratory at the age of 3 weeks, and the experiment was initiated after a 2-week acclimation period. The two study subjects (C04M3 = male, C13F5 = female) were housed separately along with four other animals in 840 cm^2^ polycarbonate cages (Unno Type III, 38.2 x 22.0 cm) consisting of dust-free, 2-5 mm aspen bedding (ssniff, Germany), aspen nesting material, mixed sized chewing sticks, a small enrichment tube and *ad libitum* access to water. Mice were fed once per day a total of 5 g of purified chow per animal. Cages were maintained in HPP750Life (Memmert, Germany) climate chambers that enabled full control and tracking of abiotic conditions. Simulated light cycles consisted of 10.5 hours of daylight and darkness and 3 hours of transitioning twilight time. Temperature was maintained at 24°C, humidity at 70%, and ventilation at 28 renovations/hour. Faecal samples were collected once a week for a period of 11 weeks. The animals were isolated in sterile cages for 30 minutes and 80 mg of faecal material per animal was collected at each sampling point. Samples were immediately stored in 1 ml of DNA/RNA Shield presentation buffer (Zymo) and frozen at -20°C until DNA extraction.

### Data generation

Faecal samples (n=22) were extracted using the in-house developed DREX protocol (details in Bozzi et al. (19)), preceded by a 10-minute bead-beating step at 30 Hz in 2 ml e-matrix tubes (MP Biomedical, USA) using a Tissuelyser II (Qiagen). Molarity and fragment-length distribution of the extracts were measured using a Tapestation. Due to the DNA amount requirements and data generation costs, two different strategies were employed to generate short- and long-read sequencing data (Fig. 1). For short-read sequencing, 200 ng of each extract were fragmented using a Covaris LE220R focused-ultrasonicator to 320-420 bp-long DNA fragments, and 11 individual sequencing libraries per animal were prepared using the BEST protocol (20). DNA sequencing was performed on an Illumina NovaSeq 6000 platform, using S4 150 paired-end chemistry, and aiming for ca. 4 Gbp (1 giga base-pair = 1×10^9^ bp) of data per library (ca. 40 Gbp per animal). For long-read sequencing, the 11 DNA extracts (145.5 ng each) of each individual were pooled to prepare a single library. Fragmented DNA (∼7,000 bp) was prepared for PacBio sequencing using the SMRTbell® express template prep kit 2.0 (Pacific Bioscience). SMRTbell® libraries were bound with sequencing primer v5 and Sequel II DNA Polymerase 2.0 using Sequel® II Binding Kit 2.2. Bound complexes were sequenced on a PacBio Sequel IIe platform using Sequencing Reagents plate 2.0. PacBio Circular Consensus Sequence (CCS) reads are produced by calling consensus of subreads of a single DNA molecule ligated with hairpin adapters, resulting in a circularised DNA molecule and allowing several passes of DNA polymerase on each strand. Circular consensus reads were generated onboard and only HiFi reads (Phred score >Q20) were used for downstream analysis. Each individual pool yielded approximately 20 Gbp of HiFi reads per library, with an average read size of around 7 Kbp.

**Figure 1.**
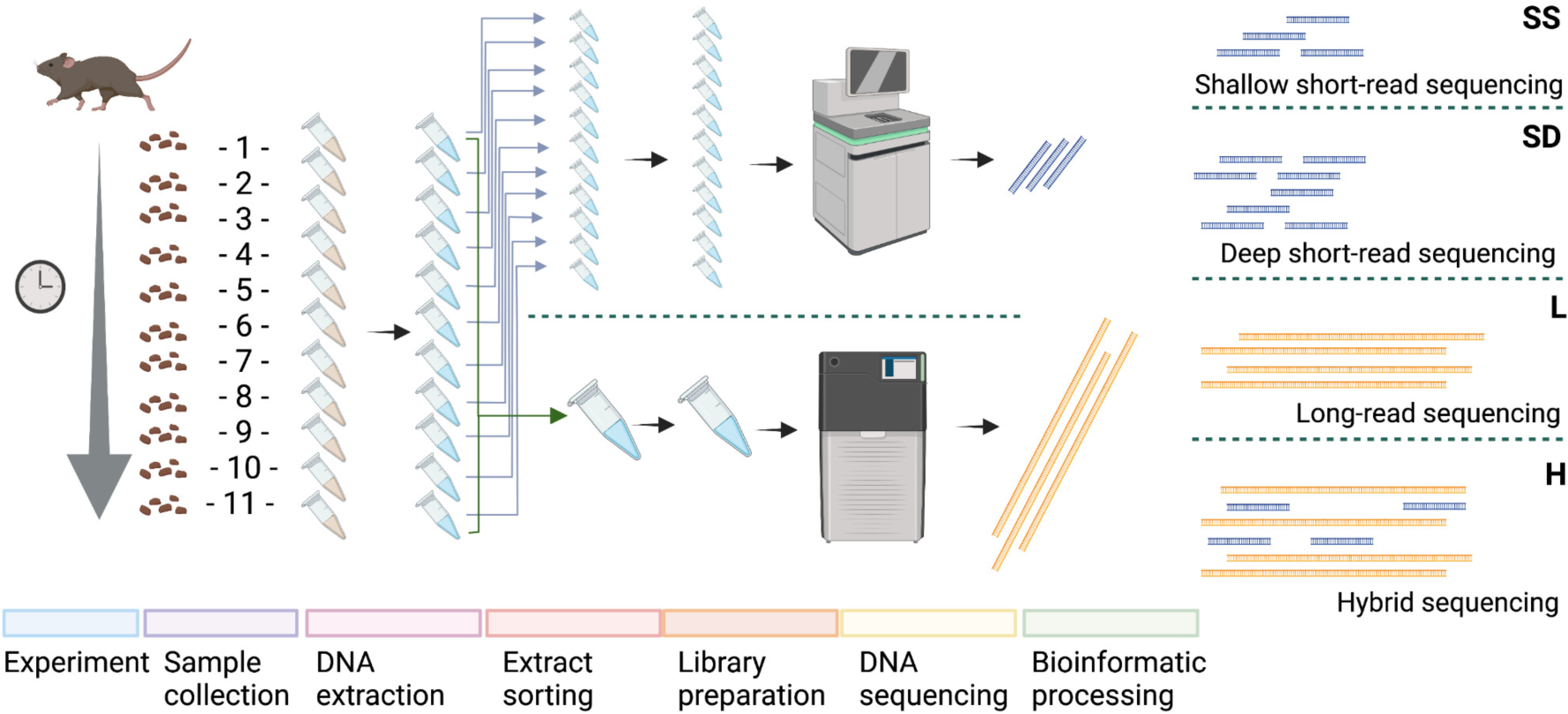
Overview of study design. Faecal samples (n=22), were collected from two mice across a longitudinal study and processed using two different laboratory protocols to generate short-read and long-read sequencing data, resulting in four datasets compared in the study. SS = short read 20 Gbp, SD = short read 40 Gbp, L = long read 20 Gbp, H = hybrid assembly 40 Gbp.

### Data analysis

Raw data generated from Illumina (n=22) and PacBio (n=2) were processed following four contrasting strategies: SR20) short-read assembly from 20 Gbp/mouse of data, SR40) short-read assembly from 40 Gbp/mouse of data, LR) long-read assembly 20 Gbp/mouse, and HY) hybrid assembly (20 Gbp of short reads + 20 Gbp of long reads per individual). Raw short-read sequencing data was preprocessed using the Earth Hologenome Initiative (EHI) preprocessing snakemake pipeline (https://github.com/earthhologenome/EHI_bioinformatics), briefly: Fastp v0.23.1 (21) was used to trim adapters and low-quality bases, following by mapping using Bowtie2 v2.4.4 (22) and samtools v1.12 (23) to the *Mus musculus* reference genome (GRCm39) to remove host reads. The filtered reads were then randomly subsampled using seqtk v1.3 (seed = 1337) (https://github.com/lh3/seqtk).

Short-read coassemblies (SR20 & SR40) were constructed using the EHI snakemake pipeline (https://github.com/earthhologenome/EHI_bioinformatics), briefly: metaSpades v3.15.3 was used for assembly (-k 21,33,55,77,99), followed by the MetaWRAP (24) binning and refinement pipelines, which use MaxBin2 (25), MetaBAT2 (26), CONCOCT (27), and CheckM (28). For the hybrid assembly (HY), metaSpades v3.15.3 (29) was used with the--pacbio flag. Binning and refinement were performed as above. Long reads (LR) were assembled using hifiasm-meta r63 (30), and binning/refinement was performed using the Pacific Biosciences HiFi-MAG-Pipeline (https://github.com/PacificBiosciences/pb-metagenomics-tools) (release 2.0.2), which uses CheckM 2 (31), semibin (32), and DAS_tool (33). QUAST v5.2.0 (34) was used to generate assembly metrics, and CoverM 0.6.1 (https://github.com/wwood/CoverM) was used to calculate the percentage of reads mapping to coassemblies. GTDB-tk (v2.1.0; database = R207_v2) was used to taxonomically annotate MAGs (35, 36).

MAG dereplication was performed using the EHI snakemake pipeline (https://github.com/earthhologenome/EHI_bioinformatics), briefly: dRep v3.4.0 (37) was run at 98% ANI (average nucleotide identity). Dereplicated MAGs were then concatenated and indexed, before being used as a reference for Bowtie2 mapping. Final count tables were generated with CoverM. antiSMASH v6.1.1 (38) was used to annotate biosynthetic gene clusters, and Barrnap v0.9 (https://github.com/tseemann/barrnap) was used to annotate 16S rRNA genes. For each strategy, we analysed the seven metrics defined in Table 1.

**Table 1.**
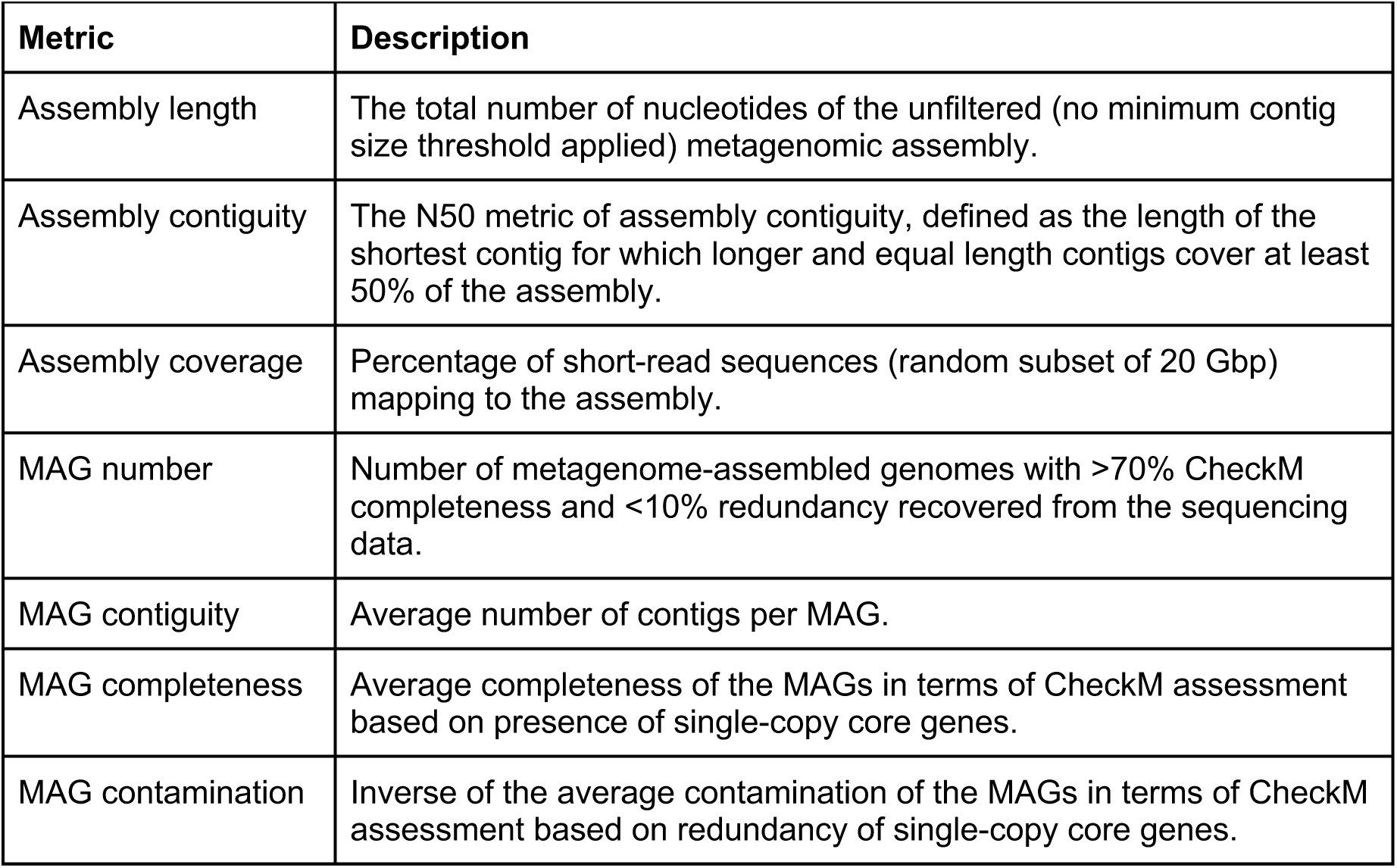
Description of the seven metrics considered for assessing the performance of the different sequencing strategies.

Complete circular sequences were identified from metagenomes assembly graphs for each dataset based on circularity using SCAPP (39) and annotated as plasmid, viral or chromosomal sequences based on specific marker genes using viralVerify v1.1 (40). Short and long reads were mapped to respective assemblies using minimap2 v2.17 (41) using “-ax sr” and “-ax map-pb” options respectively for short and long reads. Short and long reads mapped to Hybrid 40 Gbp assembly were merged before the SCAPP step. Circular contigs from metagenomes assembly graphs were extracted using SCAPP (39) using each dataset assembly and respectively mapped reads. Circular contigs were classifed as plasmid, viral or chromosomal sequences using viralVerify v1.1 (40).

Summary statistics and visualizations were performed in R using the tidyverse (42) and phyloseq (43) packages. For code specifics, see the github repository (https://github.com/EisenRa/2023_HiFi_comparison_mice). Raw sequencing reads are available at ENA accession: PRJEB65885.

### Cost estimations

Cost estimations were carried out through averaging prices obtained in early 2023 from multiple commercial sequencing service providers for DNA extraction, library preparation and sequencing short- and long-read sequencing data (Table 2).

**Table 2.** Approximate cost estimations (as for mid 2022) for generating short-read (Illumina NovaSeq 6000 S4) and long-read sequencing data (PacBio Sequel IIe HiFi), based on average quotes of multiple sequencing service providers.

|  | Cost per DNA extraction | Cost per library preparation | Sequencing cost per GB | Total cost of 20GB of data |
| --- | --- | --- | --- | --- |
| Short-read sequencing | 20 USD | 50 USD | 10 USD | 270 USD |
| Long-read sequencing | 20 USD | 350 USD | 200 USD | 4,370 USD |

## Results and Discussion

The four assembly strategies compared in this study (Fig. 1) were built from a total of 78.6 Gbp of short-read and 39.6 Gbp of long-read sequencing data generated over one NovaSeq 6000 S4 and two PacBio Sequel IIe HiFi sequencing runs, respectively. Both sequencing strategies yielded DNA reads with comparable level of quality (>40 Phred score, indicating a probability for erroneous base calling below 0.001). However, the read lengths differed significantly: the short-read sequencing yielded consistent 150 bp-long paired reads, while the long-read sequencing generated single reads with a mean length of 6,792 bp (6,351 bp for C04M3 and 7,233 bp for C13F5).

### Assembly contiguity, length, and coverage

The long-read approach yielded metagenomic assemblies with an average **assembly contiguity** 9-18 times higher than short-read and hybrid approaches (Table 3, Fig. 2/3). While the N50 value for short-read metagenomic assemblies (after filtering out <1,500 bp contigs) was around 20 Kbp (20,000 bases), the N50 values of long-read assemblies were between over 235-519 Kbp. This result is not surprising, since long-read sequences are known to improve assembly contiguity (44). The hybrid assembly exhibited a noteworthy increase in contiguity with respect to short-read only approaches (ca. 40 Kbp), but it was far lower than the values of long-read assemblies. This is likely because hybridSPAdes constructs the initial assembly graph with short-reads, using the long-reads to close gaps and resolve repeats (29). Long-read assembly also yielded the longest **assembly length** (Table 3). The hybrid assembly approach recruited the most sequences through read-mapping (96 and 97% of reads), thus exhibiting the highest **assembly coverage** across all strategies. Yet, all approaches managed to capture >93% of the metagenomic reads, indicating that most of the complexity in these samples was captured.

**Figure 2.**
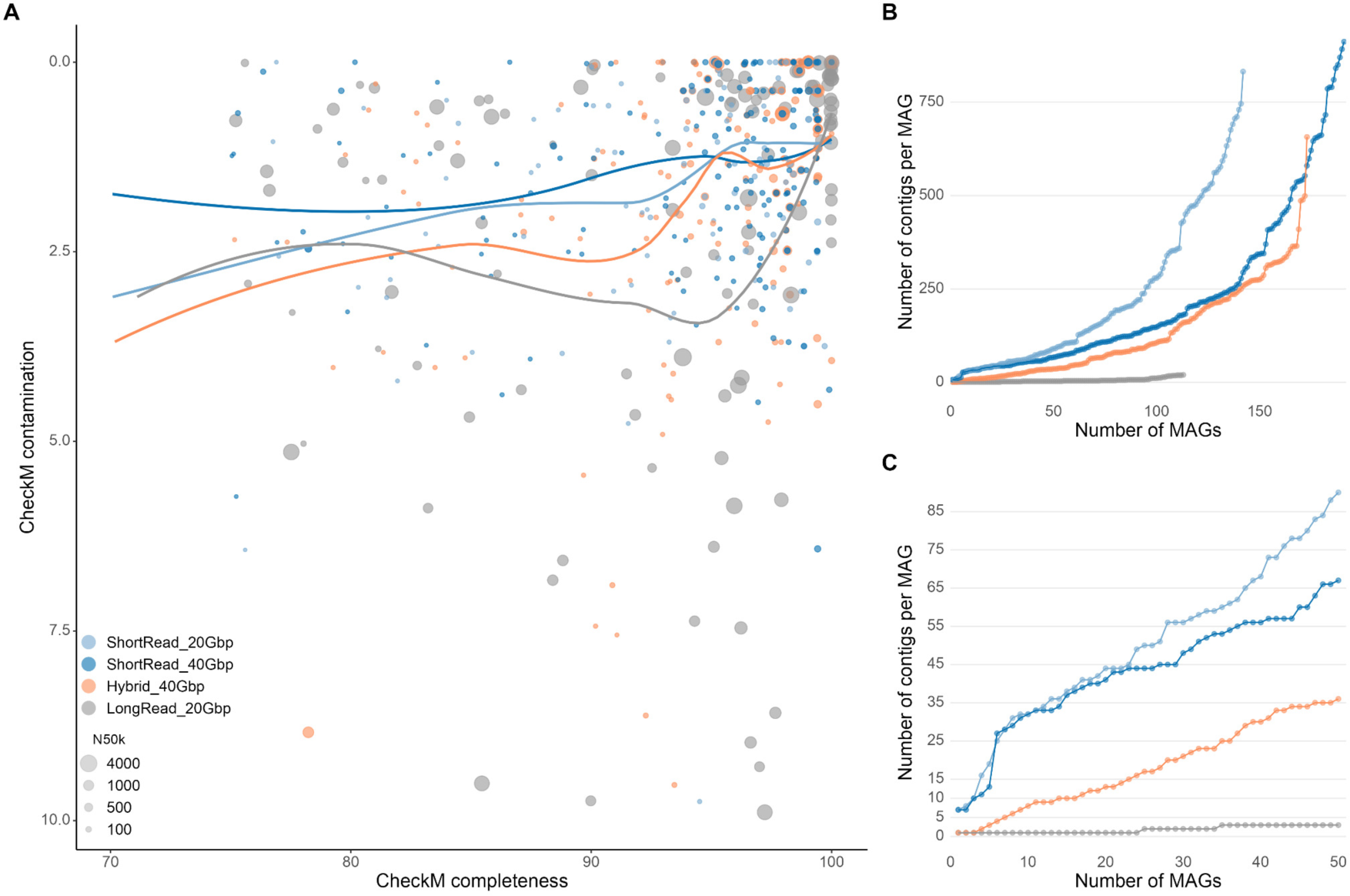
Statistics of the assemblies and MAGs recovered through the different strategies. B) all MAGs, C) top 50 most contiguous MAGs.

**Table 3.** Assembly statistics.

| Sample | Assembly length (Mbp) | Assembly contiguity (N50 Kbp) | Assembly coverage (short-reads mapped) |
| --- | --- | --- | --- |
| C04M3_Short_read_20Gbp | 345 | 20 | 93% |
| C13F5_Short_read_20Gbp | 295 | 20 | 94% |
| C04M3_Short_read_40Gbp | 451 | 30 | 95% |
| C13F5_Short_read_40Gbp | 339 | 24 | 96% |
| C04M3_Hybrid_40Gbp | 443 | 39 | **96%** |
| C13F5_Hybrid_40Gbp | 387 | 41 | **97%** |
| C04M3_Long_read_20Gbp | **505** | **236** | 94% |
| C13F5_Long_read_20Gbp | **503** | **520** | 96% |

### Quantity of recovered MAGs

The four strategies yielded a total of 619 redundant MAGs, which resulted in 125 non-redundant MAGs after dereplication at 98% ANI (Table 4, Fig. 2). Short-read approaches yielded a higher **number of MAGs** than long-read-based strategies (Fig. 3). For the same amount of nucleotide data generated (20 Gbp), the short-read approach resulted in 10 more dereplicated (at 98% ANI) MAGs compared to the long-read approach. The analysis of MAG-recovery through different strategies showed that the deficit of MAGs in the long-read approach was mainly due to the inability to recover less abundant bacteria (Fig. 4). The MAGs missed by the long-read approach accounted for on average 9% of the mapped reads. The hybrid approach recruited the highest number of reads (93% and 94%) into the dereplicated MAG catalogue. In contrast, the long-read approach recruited the lowest number of reads (62% and 70%).

**Figure 3.**
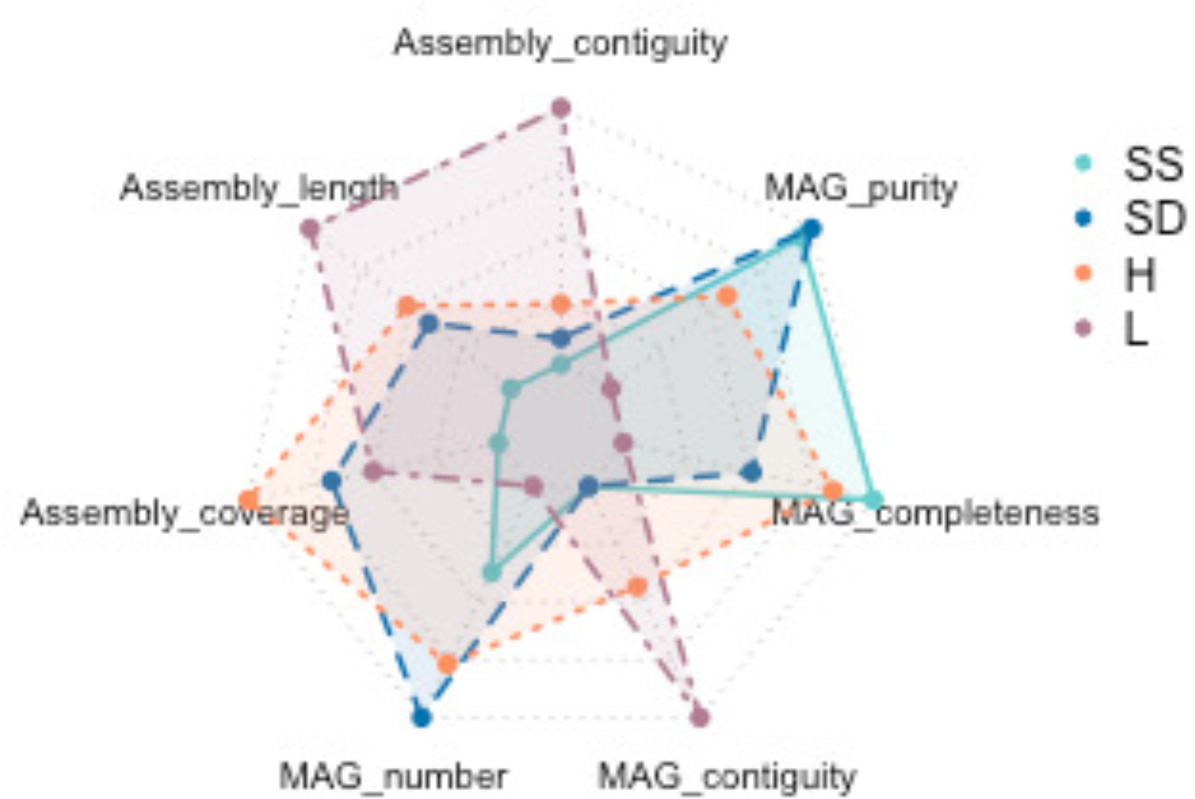
Overview of the seven metrics employed for assessing the performance of the different approaches for genome-resolved metagenomics. SS = short read 20 Gbp, SD = short read 40 Gbp, L = long read 20 Gbp, H = hybrid assembly 40 Gbp.

**Figure 4.**
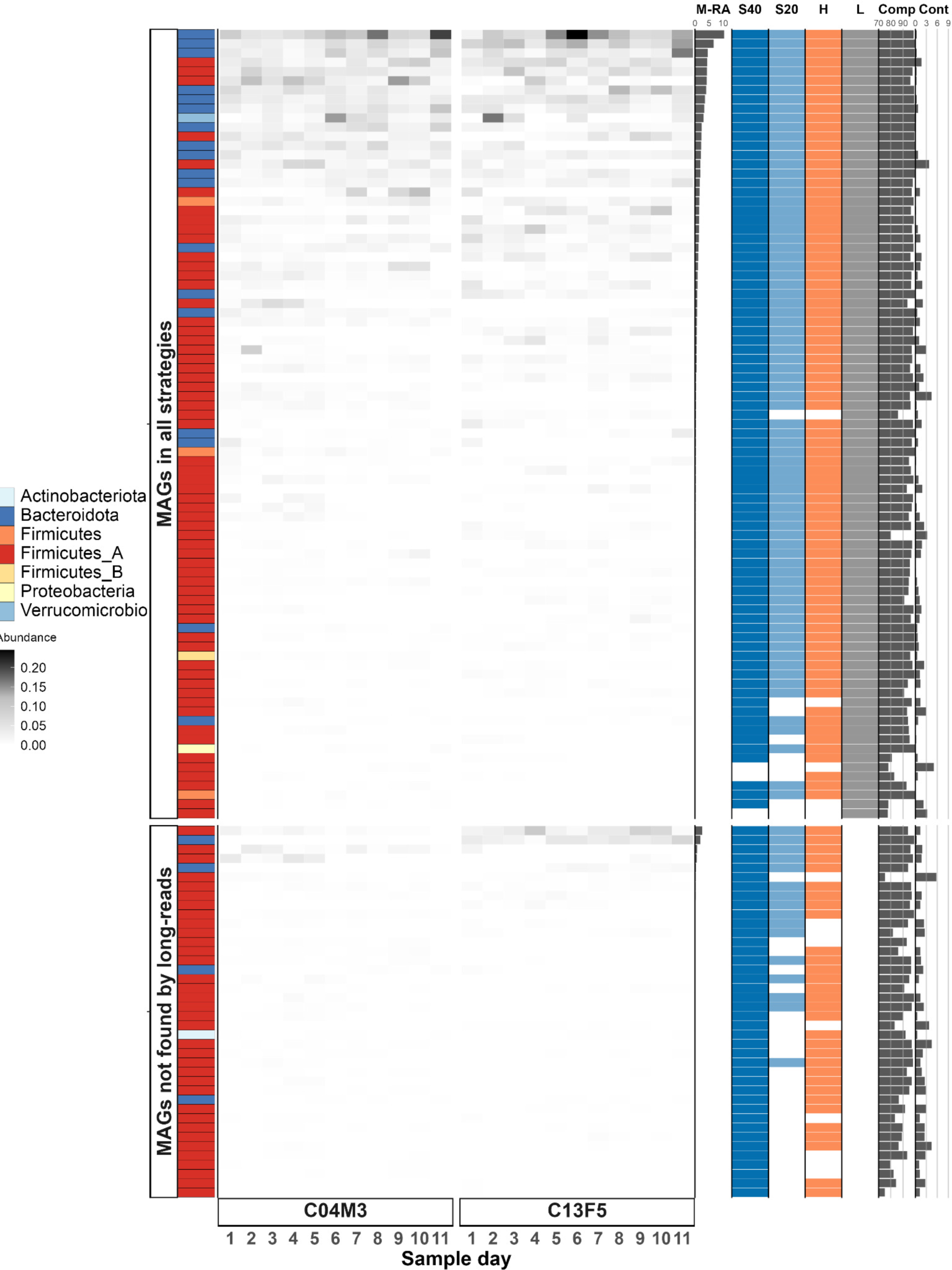
Overview of MAGs reconstruction performance by the different strategies (S40 = 40 Gbp short read, S20 = 20 Gbp short read, H = hybrid, L = HiFi long reads. All MAGs (619) were dereplicated at 98%. Each MAG is a row and each column is a sample. Filled cells indicate the relative abundance of reads (20 Gbp short reads) mapping to a given MAG. The MAGs are faceted by whether or not they were assembled using long-reads. M-RA = mean relative abundance of reads mapping to a given MAG. The four coloured columns indicate whether a given assembly strategy reconstructed a MAG. Comp and Cont = CheckM estimates for MAG completeness and contamination, respectively.

**Table 4.** MAG statistics.

| Sample | MAG coverage (short-reads mapped) | MAG number | MAG number dereplicated | MAG contiguity (mean # of contigs) | MAG completeness | MAG contamination | Circularized MAGs |
| --- | --- | --- | --- | --- | --- | --- | --- |
| C04M3_Short_read_20Gbp | 87% | 75 | 47 | 230 | 92.7% | 1.6% | 0 |
| C13F5_Short_read_20Gbp | 87% | 67 | 46 | 225 | 94.8% | 1.3% | 0 |
| C04M3_Short_read_40Gbp | 87% | **108** | 80 | 212 | 92.4% | 1.4% | 0 |
| C13F5_Short_read_40Gbp | 89% | **83** | 49 | 244 | 93.3% | 1.4% | 0 |
| C04M3_Hybrid_40Gbp | **93%** | 90 | 62 | 127 | 93.3% | 1.8% | 1 |
| C13F5_Hybrid_40Gbp | **94%** | 83 | 52 | 135 | 93.6% | 1.8% | 2 |
| C04M3_Long_read_20Gbp | 70% | 52 | 35 | **5** | 93.3% | 2.3% | **11** |
| C13F5_Long_read_20Gbp | 62% | 61 | 48 | **4** | 90.5% | 2.4% | **10** |

### Quality of recovered MAGs

Long-read sequencing outperformed the rest of the strategies in terms of MAG quality (Fig. 3). **MAG contiguity** was significantly better for long-read sequencing-based MAGs, with an average of 5 contigs per MAG versus an average of >200 contigs for MAGs reconstructed from short-read data (Table 4, Fig. 2). There was no major difference in **MAG completeness** among the assembly strategies (Table 4, SI Fig. 1). The long-read strategy successfully recovered 21 circularised genomes (14 dereplicated at 98%), of which only two had CheckM completeness scores of 100%, highlighting that SCCG-based approaches are not perfect for assessing MAG completeness (45). **MAG contamination** scores were generally low (lower scores indicate less contamination) and similar for all strategies, with the long-read approach exhibiting slightly worse values (Table 4, SI Fig. 1). Secondary metabolite biosynthetic gene clusters (BGCs) can be difficult to assemble using short-reads (46). We used antiSMASH to compare how many BGCs were reconstructed in each of the assembly approaches. While the short-read and hybrid approaches yielded the highest number of BGCs (334), when normalised by the number of MAGs in each approach, long-reads performed similarly (Table 5). Additionally, long-read assembly reconstructed over twice as many complete BGCs and substantially longer BGCs compared to the other approaches (Table 5). We did not observe marked differences in the diversity of BGCs between the different assembly approaches (SI figure 2). Finally, it’s well known that 16S rRNA genes are particularly difficult to assemble and bin using short-reads (47). We found that long-read assembly substantially improved the reconstruction of 16S rRNA genes compared to short-reads (18 versus 365) (SI figure 3).

**Table 5.** Secondary metabolite biosynthetic gene cluster (BGC) results from antiSMASH.

| Assembly | Number of BGCs | BGCs/MAGs | Complete BGCs | Mean BGC length (bp) |
| --- | --- | --- | --- | --- |
| Short_read_20Gbp | 262 | 1.85 | 36.3% | 17,511 |
| Short_read_40Gbp | **334** | 1.75 | 38.6% | 18,274 |
| Hybrid | **334** | 1.93 | 45.2% | 20,213 |
| Long_read | 219 | **1.94** | **96.3%** | **25,268** |

### Plasmids, viruses and assembled circular elements

Plasmids, viruses and other mobile genetic elements (MGEs) are of particular biological interest both in terms of public health and microbial adaptation considering that the vast majority of antibiotic resistance acquisitions in pathogenic bacteria are mediated by plasmids (48) and that viruses are an important, albeit overlooked, component of the gut microbiome (49). Plasmids are also central vectors for the genetic exchange between bacterial chromosomes (50). However, the study of plasmids in the past decade has been greatly hampered due to challenges associated with plasmid assembly using short-read sequencing (51). A definite advantage of long-in contrast to short-reads is the ability to span repeats of a few kilobases within a single sequencing read, and to capture entire circular plasmids within a single read— increasing confidence that the assembled plasmids are not chimeric. To test the advantage of long-over short-reads for plasmid metagenomic, we extracted circular sequences from each metagenome assembly graph and assigned these contigs to plasmid, viral or chromosomal sequences. We counted the number of and total length of circularised sequences obtained from each type of datasets (Fig. 5; Table 6). The number of circularized contigs varies significantly between short and long reads strategies. For the two short reads and hybrid strategies, between 1510 and 1980 circular contigs were identified (Table 6) while only 123 were recovered for the long-reads only approach. Despite the order of magnitude fewer circular contigs obtained for long-reads, the total length of these reached 42.4 Mbp, while short reads and hybrid approaches yielded between 71 Mbp and 89.2 Mbp, indicating that circular sequences obtained by long-reads assemblies are significantly larger than with short-reads.

**Figure 5.**
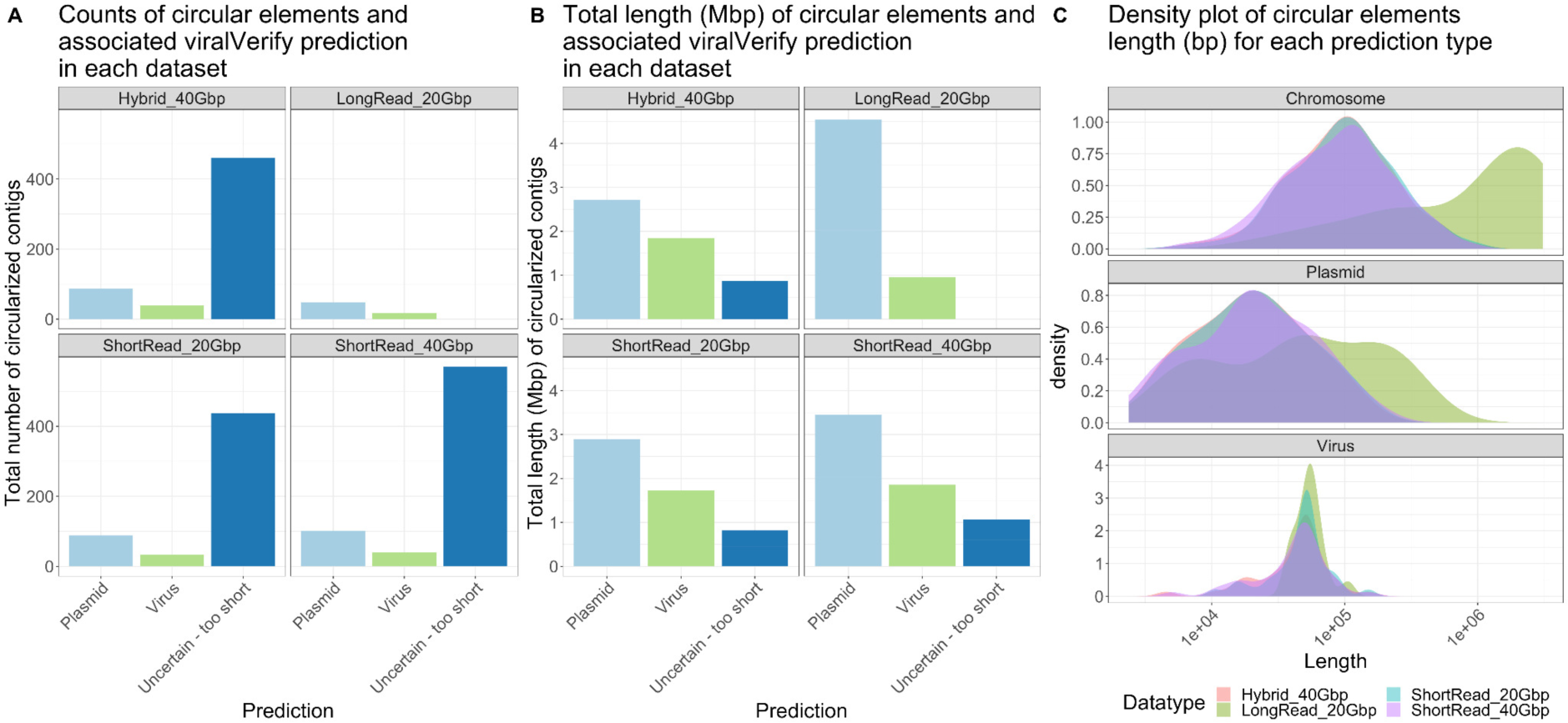
Summary of circular contigs extraction from each dataset type annotated as plasmid, virus or too short to be confidently assigned. A) Total number of circular contigs identified and annotated by viralVerify. B) Total length in megabase pairs (Mbp) of circular contigs. C) Circular contig length density for Chromsome, Plasmid and Virus prediction by viralVerify. Counts and total length of all predictions are displayed in Figure S3.

**Table 6.** Plasmids and circular contigs statistics.

| Assembly | Number of circular contigs | Total length of circular contigs (bp) | Plasmid* total length (bp) | Virus* total length (bp) | Chromosome* total length (bp) | Uncertain* total length (bp) |
| --- | --- | --- | --- | --- | --- | --- |
| Short_read_20Gbp | 1510 | 7.10+07 | 2.89E+06 (4.07%) | **1.73E+06 (2.43%)** | 6.07E+07 (85.48%) | 5.7E+06 (8.02%) |
| Short_read_40Gbp | **1980** | **8.92E+07** | 3.45E+06 (3.87%) | 1.86E+06 (2.09%) | 7.53E+07 (84.43%) | 8.57E+06 (9.61%) |
| Hybrid | 1624 | 7.77E+07 | 2.72E+06 (3.5%) | 1.85E+06 (2.38%) | **6.67E+07 (85.87%)** | 6.41E+06 (8.26%) |
| Long_read | 123 | 4.24E+07 | **4.54E+06 (10.72%)** | 9.52E+05 (2.25%) | 3.6E+07 (84.97%) | **8.75E+05 (2.07%)** |

\* based on viralVerify prediction, see methods.

A number of genetic structures on chromosomes contain repeats, for example, transposons and insertion sequences possess inverted repeats at their ends (52). Mobile genetic elements such as integrons or transposons will make circular double stranded DNA intermediates when excised from the chromosome during transposition and can be part of the total extracted DNA pool. Repeats and circular chromosomal structures will produce a circular path in contig assembly graphs and will be extracted as circular contigs by SCAPP. In addition, many dsDNA viruses have circular genomes. Circular contigs were therefore classified using marker genes specific for chromosomal, plasmid or viral sequences with viralVerify (40) (Fig. 5, SI figure 3). Most circular contigs obtained with short reads and hybrid approaches were too short to be classified as chromosomal or plasmids sequences (Fig. 5A, SI figure 3A). In contrast, most circular sequences from the long-read approach were annotated as plasmids, with no sequences too short for classification. When comparing predictions of replicon type, all approaches showed that chromosomal contigs account for the majority (84.43%-85.87%) of total circular contigs length (Table 6). Both short-reads approaches, and hybrid yielded between 9.6% and 8.6% of uncertain sequence and only between 4%-2.9% of plasmid sequences while for the long-read approach only 2% of the contigs could not be assigned and up to 10.7% of plasmid circular sequences were assembled from 20Gb of PacBio data. This indicates that circular sequences obtained by long-read approach were easier to classify and recovered more larger plasmid sequences above 100 Kb which were not recovered in short-reads datasets (Fig. 5C).

Short- and long-read approaches performed equally well to recover circular viral genomes. All methods were in agreements regarding their size distribution—centered between 25Kbp and 75Kbp (Figure 6C). This is in accordance with known sizes of DNA viruses infecting Bacteria and Archaea that are likely the most abundant hosts in the gut microbiome obtained from fecal samples (53–55). This suggests that if viruses are the targeted diversity for a study, the addition of long-reads may not substantially improve results. In contrast, if plasmids are of focus, long-reads can recover sizes that fit the broad distribution of known plasmids from databases (56). However, larger size plasmids (>100 Kb) are notably absent from short-reads datasets and could only be found with the long-read approach in the same samples. This is an important consideration when investigating the microbial ecology of plasmids where large conjugative plasmids play a prominent role (57).

## Conclusions

In conclusion, our study compared the performance of different sequencing approaches for shotgun metagenomics using faecal DNA extracts from lab mice. By assessing seven performance metrics across four combinations of sequencing depth and technology, we gained valuable insights into the strengths and limitations of the individual approaches. Our results reveal a tradeoff between the use of short-read and long-read sequencing for genome-resolved metagenomics. While deep long-read sequencing produced the highest-quality assemblies, the associated costs remain prohibitive. The optimal sequencing strategy depends on the specific goals and priorities of the study. Researchers need to carefully evaluate the balance between the quantity and quality of recovered genomes, and extrachromosomal elements considering available resources and project requirements. There is no one-size-fits-all approach, and a thoughtful and tailored selection of sequencing strategies is essential. Additionally, our study contributes new HiFi long-read sequencing data for the extensively studied laboratory mouse, which will serve as a valuable resource for future benchmarks and bioinformatic developments.

## Supplementary figures

**SI figure 1.**
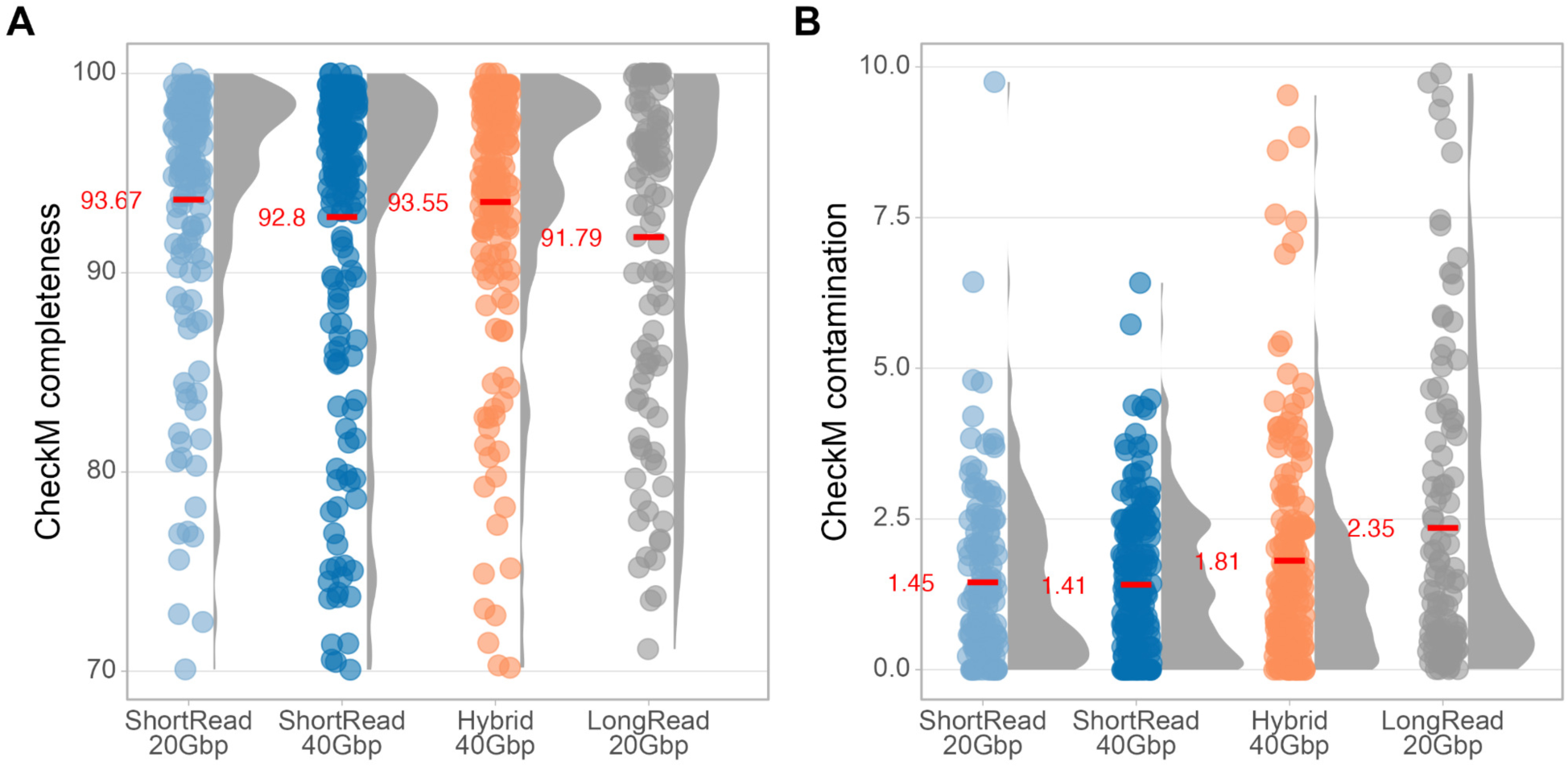
CheckM completeness (A) and contamination (B) scores for the metagenome assembled genomes.

**SI figure 2.**
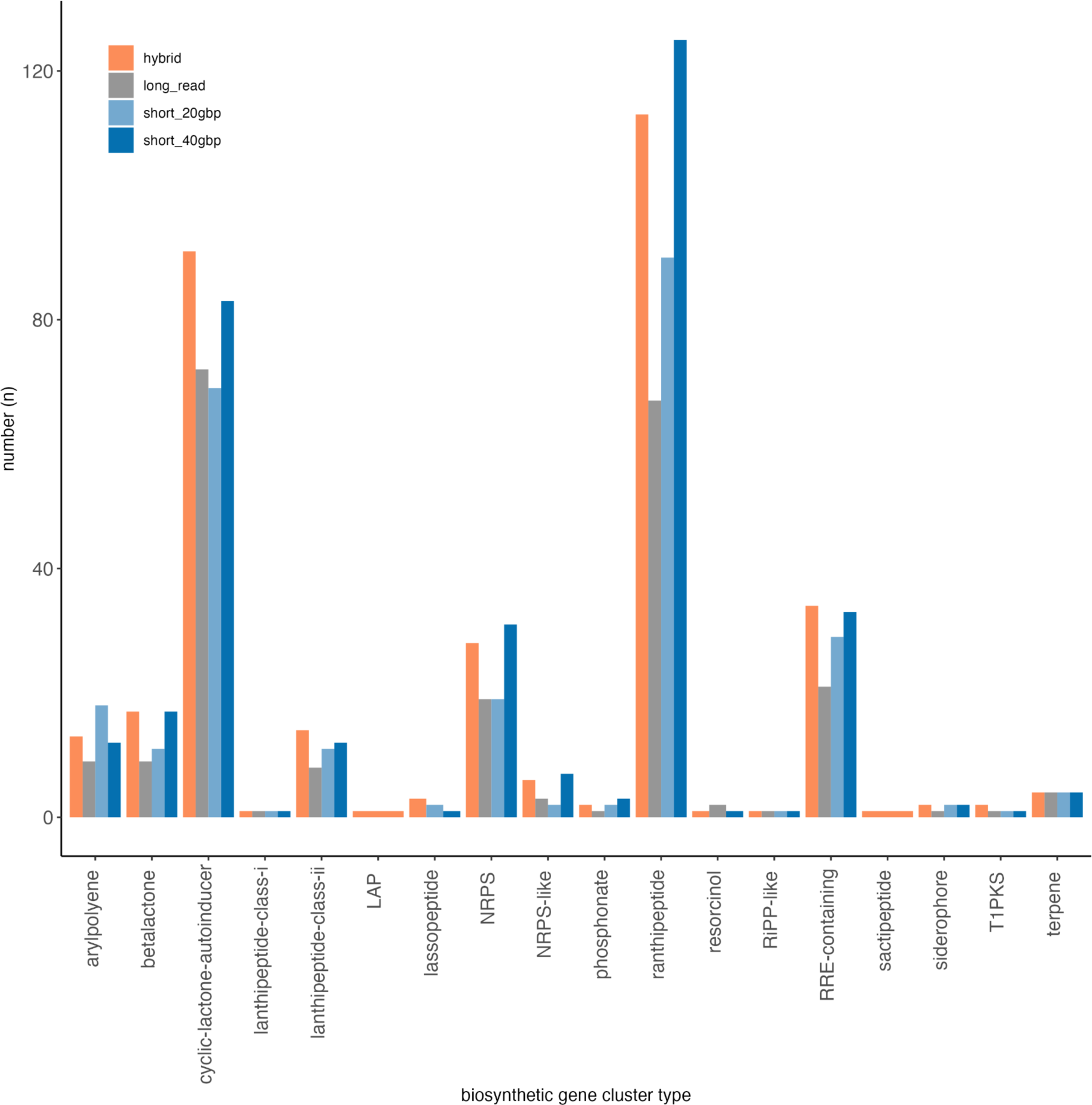
Counts of different biosynthetic gene clusters predicted by antiSMASH.

**SI figure 3.**
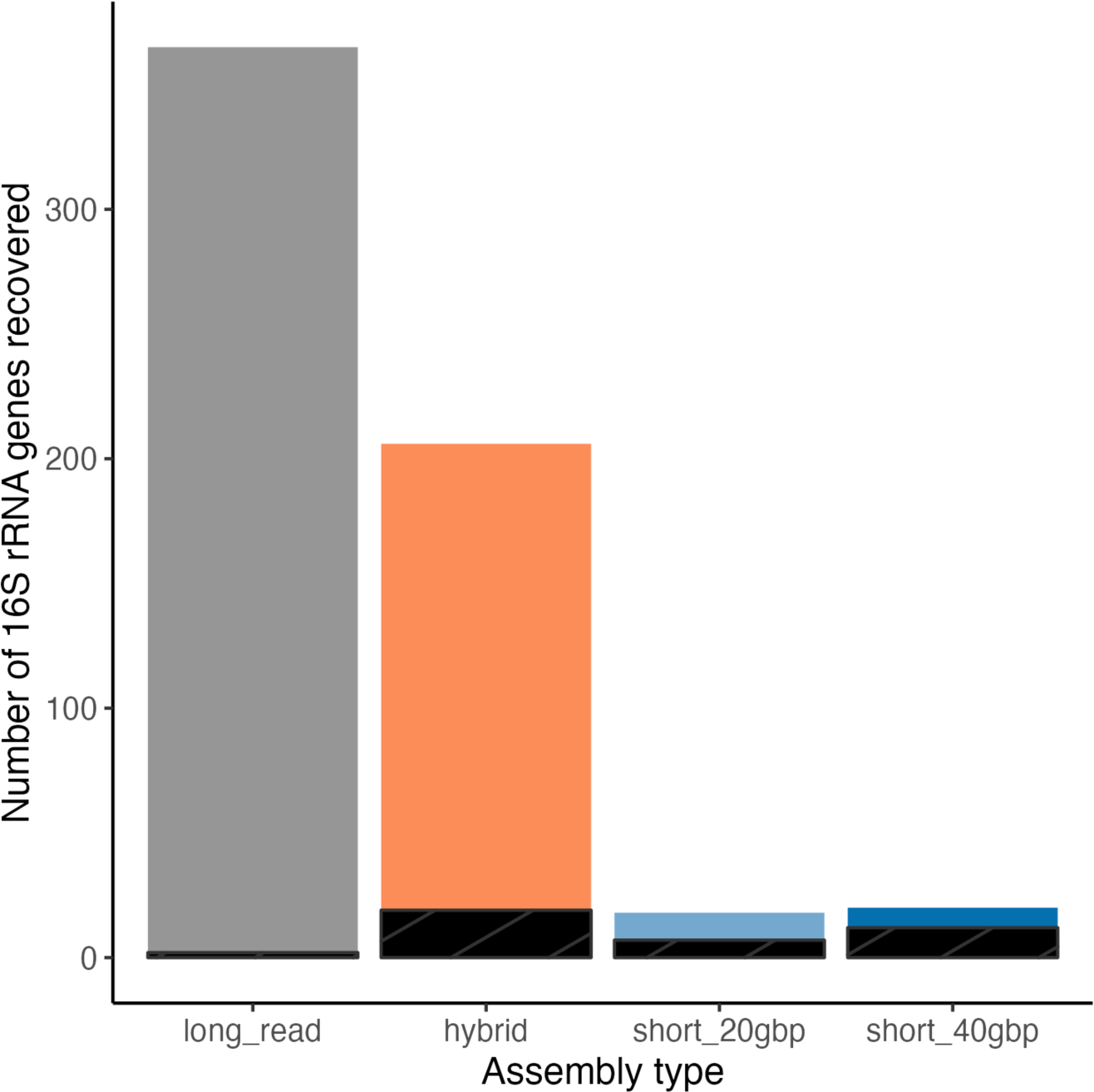
Number of 16S rRNA genes recovered by different assembly types. Black bars indicate incomplete 16S rRNA genes (<1,450 bp length).

**SI figure 4.**
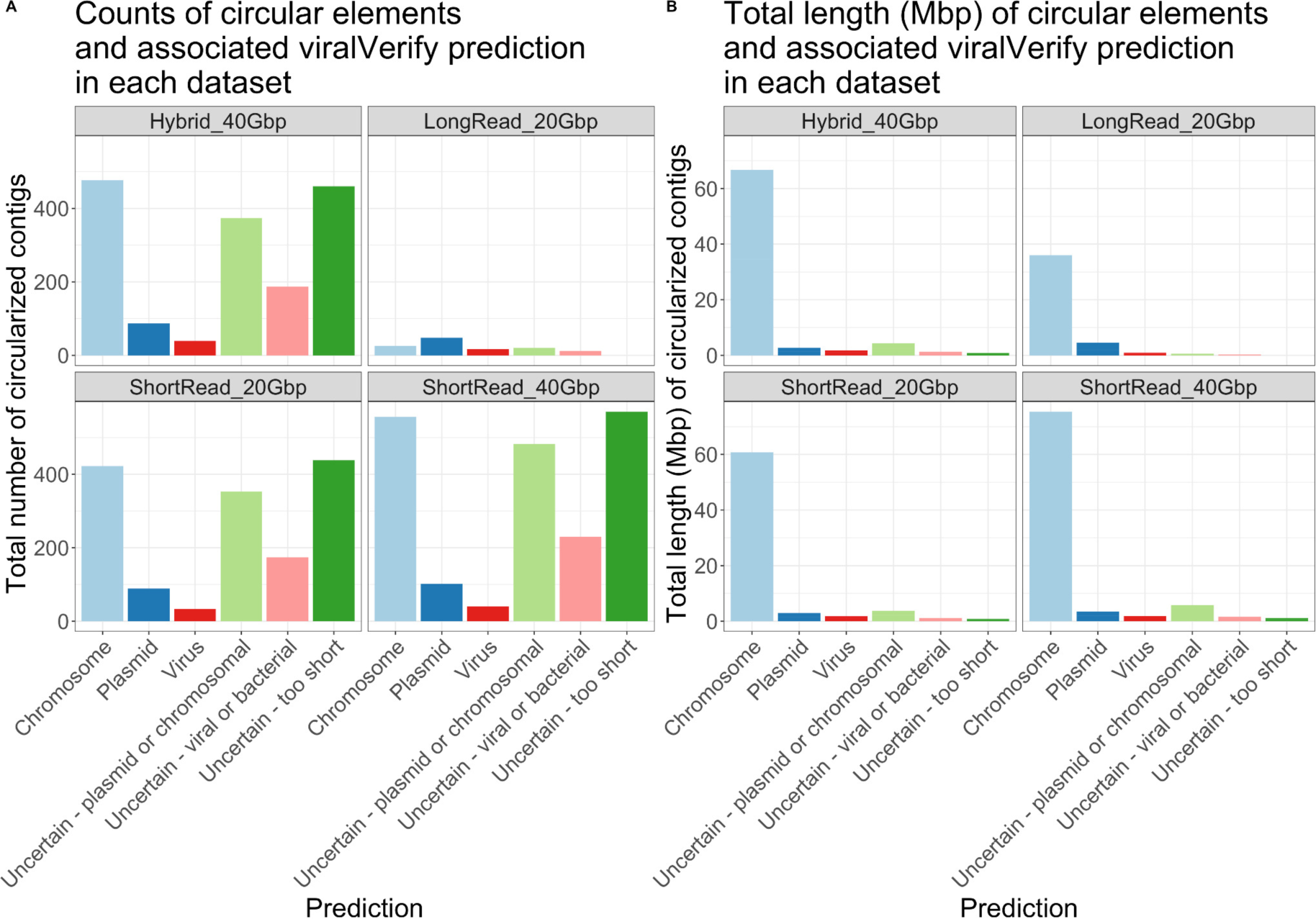
Summary of circular contigs extraction from each dataset type annotated by viralVerify. A) Total number of circular contigs identified. B) Total length in megabase pairs (Mbp) of circular contigs.

## References

1. Quince C, Walker AW, Simpson JT, Loman NJ, Segata N. 2017. Shotgun metagenomics, from sampling to analysis. Nat Biotechnol 35:833–844.

2. Almeida A, Nayfach S, Boland M, Strozzi F, Beracochea M, Shi ZJ, Pollard KS, Sakharova E, Parks DH, Hugenholtz P, Segata N, Kyrpides NC, Finn RD. 2020. A unified catalog of 204,938 reference genomes from the human gut microbiome. Nat Biotechnol 10.1038/s41587-020-0603-3.

3. Parks DH, Rinke C, Chuvochina M, Chaumeil P-A, Woodcroft BJ, Evans PN, Hugenholtz P, Tyson GW. 2017. Recovery of nearly 8,000 metagenome-assembled genomes substantially expands the tree of life. Nat Microbiol 2:1533–1542.

4. Pasolli E, Asnicar F, Manara S, Zolfo M, Karcher N, Armanini F, Beghini F, Manghi P, Tett A, Ghensi P, Collado MC, Rice BL, DuLong C, Morgan XC, Golden CD, Quince C, Huttenhower C, Segata N. 2019. Extensive Unexplored Human Microbiome Diversity Revealed by Over 150,000 Genomes from Metagenomes Spanning Age, Geography, and Lifestyle. Cell 176:649–662.e20.

5. Tyson GW, Chapman J, Hugenholtz P, Allen EE, Ram RJ, Richardson PM, Solovyev VV, Rubin EM, Rokhsar DS, Banfield JF. 2004. Community structure and metabolism through reconstruction of microbial genomes from the environment. Nature 428:37–43.

6. Shaffer M, Borton MA, McGivern BB, Zayed AA, La Rosa SL, Solden LM, Liu P, Narrowe AB, Rodríguez-Ramos J, Bolduc B, Gazitúa MC, Daly RA, Smith GJ, Vik DR, Pope PB, Sullivan MB, Roux S, Wrighton KC. 2020. DRAM for distilling microbial metabolism to automate the curation of microbiome function. Nucleic Acids Res 48:8883–8900.

7. Amarasinghe SL, Su S, Dong X, Zappia L, Ritchie ME, Gouil Q. 2020. Opportunities and challenges in long-read sequencing data analysis. Genome Biol 21:30.

8. Alkan C, Sajjadian S, Eichler EE. 2011. Limitations of next-generation genome sequence assembly. Nat Methods 8:61–65.

9. Logsdon GA, Vollger MR, Eichler EE. 2020. Long-read human genome sequencing and its applications. Nat Rev Genet 21:597–614.

10. Kim CY, Ma J, Lee I. 2022. HiFi metagenomic sequencing enables assembly of accurate and complete genomes from human gut microbiota. Nat Commun 13:6367.

11. Bickhart DM, Kolmogorov M, Tseng E, Portik DM, Korobeynikov A, Tolstoganov I, Uritskiy G, Liachko I, Sullivan ST, Shin SB, Zorea A, Andreu VP, Panke-Buisse K, Medema MH, Mizrahi I, Pevzner PA, Smith TPL. 2022. Generating lineage-resolved, complete metagenome-assembled genomes from complex microbial communities. Nat Biotechnol 40:711–719.

12. Zhang Y, Jiang F, Yang B, Wang S, Wang H, Wang A, Xu D, Fan W. 2022. Improved microbial genomes and gene catalog of the chicken gut from metagenomic sequencing of high-fidelity long reads. Gigascience 11.

13. Jiang F, Li Q, Wang S, Shen T, Wang H, Wang A, Xu D, Yuan L, Lei L, Chen R, Yang B, Deng Y, Fan W. 2022. Recovery of metagenome-assembled microbial genomes from a full-scale biogas plant of food waste by pacific biosciences high-fidelity sequencing. Front Microbiol 13:1095497.

14. Priest T, Orellana LH, Huettel B, Fuchs BM, Amann R. 2021. Microbial metagenome-assembled genomes of the Fram Strait from short and long read sequencing platforms. PeerJ 9:e11721.

15. Tao Y, Xun F, Zhao C, Mao Z, Li B, Xing P, Wu QL. 2023. Improved Assembly of Metagenome-Assembled Genomes and Viruses in Tibetan Saline Lake Sediment by HiFi Metagenomic Sequencing. Microbiol Spectr 11:e0332822.

16. Bertrand D, Shaw J, Kalathiyappan M, Ng AHQ, Kumar MS, Li C, Dvornicic M, Soldo JP, Koh JY, Tong C, Ng OT, Barkham T, Young B, Marimuthu K, Chng KR, Sikic M, Nagarajan N. 2019. Hybrid metagenomic assembly enables high-resolution analysis of resistance determinants and mobile elements in human microbiomes. Nat Biotechnol 37:937–944.

17. Chen L, Zhao N, Cao J, Liu X, Xu J, Ma Y, Yu Y, Zhang X, Zhang W, Guan X, Yu X, Liu Z, Fan Y, Wang Y, Liang F, Wang D, Zhao L, Song M, Wang J. 2022. Short- and long-read metagenomics expand individualized structural variations in gut microbiomes. Nat Commun 13:3175.

18. Jin H, You L, Zhao F, Li S, Ma T, Kwok L-Y, Xu H, Sun Z. 2022. Hybrid, ultra-deep metagenomic sequencing enables genomic and functional characterization of low-abundance species in the human gut microbiome. Gut Microbes 14:2021790.

19. Bozzi D, Rasmussen JA, Carøe C, Sveier H, Nordøy K, Gilbert MTP, Limborg MT. 2021. Salmon gut microbiota correlates with disease infection status: potential for monitoring health in farmed animals. Anim Microbiome 3:30.

20. Carøe C, Gopalakrishnan S, Vinner L, Mak SST, Sinding M-HS, Samaniego JA, Wales N, Sicheritz-Pontén T, Gilbert MTP. 2017. Single-tube library preparation for degraded DNA. Methods in Ecology and Evolution.

21. Chen S, Zhou Y, Chen Y, Gu J. 2018. fastp: an ultra-fast all-in-one FASTQ preprocessor. Bioinformatics 34:i884–i890.

22. Langmead B, Salzberg SL. 2012. Fast gapped-read alignment with Bowtie 2. Nat Methods 9:357–359.

23. Li H, Handsaker B, Wysoker A, Fennell T, Ruan J, Homer N, Marth G, Abecasis G, Durbin R, 1000 Genome Project Data Processing Subgroup. 2009. The Sequence Alignment/Map format and SAMtools. Bioinformatics 25:2078–2079.

24. Uritskiy GV, DiRuggiero J, Taylor J. 2018. MetaWRAP-a flexible pipeline for genome-resolved metagenomic data analysis. Microbiome 6:158.

25. Wu Y-W, Simmons BA, Singer SW. 2016. MaxBin 2.0: an automated binning algorithm to recover genomes from multiple metagenomic datasets. Bioinformatics 32:605–607.

26. Kang DD, Li F, Kirton E, Thomas A, Egan R, An H, Wang Z. 2019. MetaBAT 2: an adaptive binning algorithm for robust and efficient genome reconstruction from metagenome assemblies. PeerJ 7:e7359.

27. Alneberg J, Bjarnason BS, de Bruijn I, Schirmer M, Quick J, Ijaz UZ, Lahti L, Loman NJ, Andersson AF, Quince C. 2014. Binning metagenomic contigs by coverage and composition. Nat Methods 11:1144–1146.

28. Parks DH, Imelfort M, Skennerton CT, Hugenholtz P, Tyson GW. 2015. CheckM: assessing the quality of microbial genomes recovered from isolates, single cells, and metagenomes. Genome Res 25:1043–1055.

29. Antipov D, Korobeynikov A, McLean JS, Pevzner PA. 2016. hybridSPAdes: an algorithm for hybrid assembly of short and long reads. Bioinformatics 32:1009–1015.

30. Feng X, Cheng H, Portik D, Li H. 2022. Metagenome assembly of high-fidelity long reads with hifiasm-meta. Nat Methods 19:671–674.

31. Chklovski A, Parks DH, Woodcroft BJ, Tyson GW. 2022. CheckM2: a rapid, scalable and accurate tool for assessing microbial genome quality using machine learning. bioRxiv.

32. Pan S, Zhu C, Zhao X-M, Coelho LP. 2022. A deep siamese neural network improves metagenome-assembled genomes in microbiome datasets across different environments. Nat Commun 13:2326.

33. Sieber CMK, Probst AJ, Sharrar A, Thomas BC, Hess M, Tringe SG, Banfield JF. 2018. Recovery of genomes from metagenomes via a dereplication, aggregation and scoring strategy. Nat Microbiol 3:836–843.

34. Mikheenko A, Prjibelski A, Saveliev V, Antipov D, Gurevich A. 2018. Versatile genome assembly evaluation with QUAST-LG. Bioinformatics 34:i142–i150.

35. Rinke C, Chuvochina M, Mussig AJ, Chaumeil P-A, Davín AA, Waite DW, Whitman WB, Parks DH, Hugenholtz P. 2021. A standardized archaeal taxonomy for the Genome Taxonomy Database. Nat Microbiol 6:946–959.

36. Chaumeil P-A, Mussig AJ, Hugenholtz P, Parks DH. 2022. GTDB-Tk v2: memory friendly classification with the genome taxonomy database. Bioinformatics 38:5315–5316.

37. Olm MR, Brown CT, Brooks B, Banfield JF. 2017. dRep: a tool for fast and accurate genomic comparisons that enables improved genome recovery from metagenomes through de-replication. ISME J 11:2864–2868.

38. Blin K, Shaw S, Kloosterman AM, Charlop-Powers Z, van Wezel GP, Medema MH, Weber T. 2021. antiSMASH 6.0: improving cluster detection and comparison capabilities. Nucleic Acids Res 49:W29–W35.

39. Pellow D, Zorea A, Probst M, Furman O, Segal A, Mizrahi I, Shamir R. 2021. SCAPP: an algorithm for improved plasmid assembly in metagenomes. Microbiome 9:144.

40. Antipov D, Raiko M, Lapidus A, Pevzner PA. 2020. Metaviral SPAdes: assembly of viruses from metagenomic data. Bioinformatics 36:4126–4129.

41. Li H. 2018. Minimap2: pairwise alignment for nucleotide sequences. Bioinformatics 34:3094–3100.

42. Wickham H, Averick M, Bryan J, Chang W, McGowan L, François R, Grolemund G, Hayes A, Henry L, Hester J, Kuhn M, Pedersen T, Miller E, Bache S, Müller K, Ooms J, Robinson D, Seidel D, Spinu V, Takahashi K, Vaughan D, Wilke C, Woo K, Yutani H. 2019. Welcome to the tidyverse. J Open Source Softw 4:1686.

43. McMurdie PJ, Holmes S. 2013. phyloseq: an R package for reproducible interactive analysis and graphics of microbiome census data. PLoS One 8:e61217.

44. Koren S, Phillippy AM. 2015. One chromosome, one contig: complete microbial genomes from long-read sequencing and assembly. Curr Opin Microbiol 23:110–120.

45. Chen L-X, Anantharaman K, Shaiber A, Eren AM, Banfield JF. 2020. Accurate and complete genomes from metagenomes. Genome Res 30:315–333.

46. Goldstein S, Beka L, Graf J, Klassen JL. 2019. Evaluation of strategies for the assembly of diverse bacterial genomes using MinION long-read sequencing. BMC Genomics 20:23.

47. Vollmers J, Wiegand S, Kaster A-K. 2017. Comparing and Evaluating Metagenome Assembly Tools from a Microbiologist’s Perspective - Not Only Size Matters! PLoS One 12:e0169662.

48. San Millan A. 2018. Evolution of Plasmid-Mediated Antibiotic Resistance in the Clinical Context. Trends Microbiol 26:978–985.

49. Cao Z, Sugimura N, Burgermeister E, Ebert MP, Zuo T, Lan P. 2022. The gut virome: A new microbiome component in health and disease. EBioMedicine 81:104113.

50. Halary S, Leigh JW, Cheaib B, Lopez P, Bapteste E. 2010. Network analyses structure genetic diversity in independent genetic worlds. Proc Natl Acad Sci U S A 107:127–132.

51. Arredondo-Alonso S, Willems RJ, van Schaik W, Schürch AC. 2017. On the (im)possibility of reconstructing plasmids from whole-genome short-read sequencing data. Microb Genom 3:e000128.

52. Tørresen OK, Star B, Mier P, Andrade-Navarro MA, Bateman A, Jarnot P, Gruca A, Grynberg M, Kajava AV, Promponas VJ, Anisimova M, Jakobsen KS, Linke D. 2019. Tandem repeats lead to sequence assembly errors and impose multi-level challenges for genome and protein databases. Nucleic Acids Res 47:10994–11006.

53. Chaudhari HV, Inamdar MM, Kondabagil K. 2021. Scaling relation between genome length and particle size of viruses provides insights into viral life history. iScience 24:102452.

54. Liang G, Bushman FD. 2021. The human virome: assembly, composition and host interactions. Nat Rev Microbiol 19:514–527.

55. Fiege JK, Langlois RA. 2022. Embracing the heterogeneity of natural viruses in mouse studies. J Gen Virol 103.

56. Smillie Chris, Garcillán-Barcia M. Pilar, Francia M. Victoria, Rocha Eduardo P. C., de la Cruz Fernando. 2010. Mobility of Plasmids. Microbiol Mol Biol Rev 74:434–452.

57. Bottery MJ. 2022. Ecological dynamics of plasmid transfer and persistence in microbial communities. Curr Opin Microbiol 68:102152.

